# A conserved sRNA regulates mucin adhesion and gut colonization across the Enterococcaceae

**DOI:** 10.1101/2025.11.10.687296

**Authors:** Sierra Bowden, Kevin C. Jennings, Soumaya Zlitni, Erin F. Brooks, Megan E. Davin, Aravind Natarajan, Leonard Asare, Brayon J. Fremin, Paul Wilmes, Robert L. Hettich, Nita H. Salzman, Ami S. Bhatt

## Abstract

Enterococci, particularly *E. faecalis*, can survive in diverse settings within and outside human hosts. The capacity of *E. faecalis* to colonize these locations relies on its ability to adapt by altering gene expression in response to environmental exposures. One mechanism for quickly altering gene expression is through regulation by small noncoding RNAs (sRNAs); sRNAs can regulate one or many target genes and either up- or down-regulate transcript stability and protein expression. While many sRNAs have been predicted in *E. faecalis,* few have experimentally established target mRNAs or physiological functions. Here, we investigate the targets, function, and mechanism of Enterococcus sRNA 84. We found that sRNA 84 is conserved within the family Enterococcaceae, suggesting that it plays a role in the regulation of core genes and functions. RNA sequencing and proteomic analysis revealed that the absence of sRNA 84 led to downregulation of many cell surface proteins, including mucin-binding proteins. Consistent with these findings, an sRNA 84 knockout strain had reduced binding to mucin *in vitro* and impaired intestinal colonization of specific-pathogen-free mice. Taken together, these data support a model whereby sRNA 84 upregulates cell surface adhesins, which subsequently facilitate host colonization through binding to mucin. sRNA 84 is one of the first sRNAs in enterococci with demonstrated targets and function. This finding establishes the conserved sRNA 84 as a potential key regulator of enterococcal host adaptation, providing insight into how these organisms adapt their gene expression to survive both within and outside animal hosts.

## Introduction

Enterococci play key roles in maintaining human health and causing disease. Emerging hundreds of millions of years ago (Lebreton et al. 2017), enterococci are an ancient member of the human gut microbiome. In particular, *Enterococcus faecalis* is present in around 80% of healthy adults, colonizing the intestines at a relative abundance below 1%. As one of the earliest colonizers of the human gut, *E. faecalis* is often present from the first week of life (Kao & Kline 2019). However, enterococci can also act as opportunistic pathogens and are a leading cause of nosocomial infections such as bacteremia, surgical wound infections, catheter-associated urinary tract infections, and endocarditis (Arias & Murray 2012). Their high levels of intrinsic and acquired antibiotic resistance, as well as their ability to survive exposure to antiseptic agents and harsh environments, makes them particularly persistent in the clinic. Vancomycin-resistant *E. faecalis* is now prevalent in hospital environments and is estimated to cause over 50,000 infections and 5,000 deaths in the United States annually (CDC 2025). While enterococcal infections are on the rise, it has in fact been known for over 100 years that enterococci are “very hardy and tenacious” (MacCallum and Hastings 1899). A hallmark of enterococci is the ability to survive in many environments; this includes various body sites (gut, blood, mouth, urinary tract), different organisms, and even outside the host on abiotic surfaces. *E. faecalis,* in particular, is a versatile generalist, and its prevalence in the gut microbiome and hospital devices may be a factor in causing many infections in immunocompromised patients.

In order to survive in these different environments, *E. faecalis* must be able to rapidly adapt to changing conditions; one way it achieves this is through regulation of gene expression. While transcript levels are largely controlled through transcription factor-based regulation, post-transcriptional regulatory mechanisms, such as small noncoding RNAs (sRNAs), rapidly adjust and fine-tune transcript and protein abundances. sRNAs are noncoding, regulatory RNAs with diverse structures, mechanisms, and functions, playing key roles in stress response and lifestyle transitions (Cao et al. 2023). One class of sRNAs is RNA-binding RNAs, which work to alter gene expression by base pairing with target mRNAs. These include antisense RNAs (which regulate the transcript encoded on the opposite strand) and intergenic *trans-*encoded sRNAs, which can regulate many targets all over the genome, through base pairing with imperfect complementarity. *Trans-*encoded sRNAs are a powerful and flexible means of gene regulation— they are energetically inexpensive, as they do not rely on expressing a protein, and can regulate many genes at once to quickly respond to environmental changes (Beisel & Storz 2010). They can also up- or downregulate targets, either by impacting mRNA stability (protecting from or promoting degradation) or protein translation (inhibiting or activating ribosome binding) (Waters & Storz 2009).

Much of what is known about sRNA mechanisms and functions was discovered in well-studied model organisms like *Escherichia coli,* which rely on a global RNA binding chaperone such as Hfq. Studying known RNA binding proteins has enabled the study of many interacting RNAs. However, many Gram-positive organisms, including *E. faecalis,* lack any known protein facilitator of sRNA-mRNA interactions (Olejniczak et al. 2022). Because of this, discovery and experimental validation of sRNAs, their targets, and functions has lagged behind. The first annotations of sRNAs in enterococci were a part of kingdom-wide computational predictions (Livny et al. 2008). Microarrays, RACE-PCR, and northern blots identified and validated new sRNAs in *E. faecalis* (Fouquier d’Hérouël et al. 2011, Shioya et al. 2011). With the advent of RNA sequencing, global annotation of RNAs in *E. faecalis* (Innocenti et al. 2015, Salze et al. 2020), *E. faecium* (Sinel et al. 2017), and *E. faecalis* and *E. faecium* (Michaux et al. 2020) greatly expanded the known sRNAs in enterococci. Grad-Seq experiments helped identify which transcripts are noncoding, and which may encode small proteins (Michaux et al. 2023). Despite the growth in discovery of new sRNAs, few have experimentally validated targets or functions (Michaux et al. 2014, Reissier et al. 2021).

In this study, we investigate the targets, function, and mechanism of Enterococcus sRNA 84. Enterococcus sRNA 84 is a *trans-*encoded RNA-binding RNA found across the family Enterococcaceae, in both *E. faecalis* and *E. faecium,* including commensal and pathogenic strains. Based on its conservation, we hypothesized that sRNA 84 plays a key role in regulating core genes, and therefore physiologically critical functions. We conducted RNA sequencing analyses and determined that sRNA 84 is important for activating many cell wall genes, likely through direct base pairing resulting in stabilization of the transcript. Without sRNA 84, transcripts encoding these cell wall adhesins are lowly abundant, thereby decreasing the ability of *E. faecalis* to bind to mucin *in vitro* and to colonize the mouse gastrointestinal tract. Regulation of cell wall adhesins enables *E. faecalis* to adapt to the dynamic environment of the human gastrointestinal tract, which may contribute to the ubiquity of *E. faecalis* as a gut commensal. sRNA 84 is one of the first sRNAs in enterococci to have demonstrated targets and function. Its rather unique conservation across the Enterococcaceae may provide insight into how these organisms adapt their gene expression to survive both in and outside of the host environment.

## Results

### Enterococci sRNA 84 is a *trans-*encoded sRNA found across the Enterococcaceae

Many sRNAs (especially RNA-binding RNAs) are thought to be species- or even strain-specific (Livny et al. 2008 and Michaux et al. 2020). However, understanding of the conservation of sRNAs across enterococci has so far been limited. Most studies on sRNAs in enterococci have been conducted in one of only a few strains, mainly the clinical isolates V583 and AUS0004 (Fouquier d’Hérouël et al. 2011, Shioya et al. 2011, Innocenti et al. 2015, Sinel et al. 2017, Michaux et al. 2020). Additionally, different nomenclature can make it difficult to compare results across studies. In order to understand the distribution and abundance of sRNAs in enterococci, we used cmscan (Nawrocki and Eddy 2013) to annotate all RNAs found in the RNA families database (Rfam) (Ontiberos-Palacios et al. 2025) across 29 representative genomes (Lebreton et. al 2017). 24 of these belong to the genus *Enterococcus,* including 2 different strains of *E. faecalis* and 3 strains of *E. faecium.* We also included genomes from other members of the family Enterococcaceae (*Melissococcus plutonius* and *Vagococcus lutrae*) as well as more distantly related species (*Lactococcus garvieae* and *Carnobacterium maltaromanticum*), and *Listeria monoytogenes* as an outgroup. We filtered for RNA families of the type ‘sRNA’ or ‘antisense’ (asRNA) to identify RNA-binding RNAs (Figure 1A). In total, we found 39 RNAs present in at least one enterococcal genome. *E. faecalis* V583 had the most, with 19 unique RNAs. This may be due to an overrepresentation of studies on V583 and on *E. faecalis* compared to the other genomes in this data set. 4 of these RNAs are unique to strain V583 and are not found in OG1RF. While many sRNAs do appear to be found in only one species, there are notable exceptions. RNAs such as RF01458, RF01470, RF01492, and RF01494 are found in many genomes, often with high copy numbers (as many as 11 for RF01492). These RNAs are all found in the outgroup *L. montocytogenes* and are distributed across a broad range of species in Rfam. Mraheil et al. posit that sRNAs such as RF01492 are located on mobile genetic elements enabling their spread to many genomes and loci (2011).

**Figure 1:**
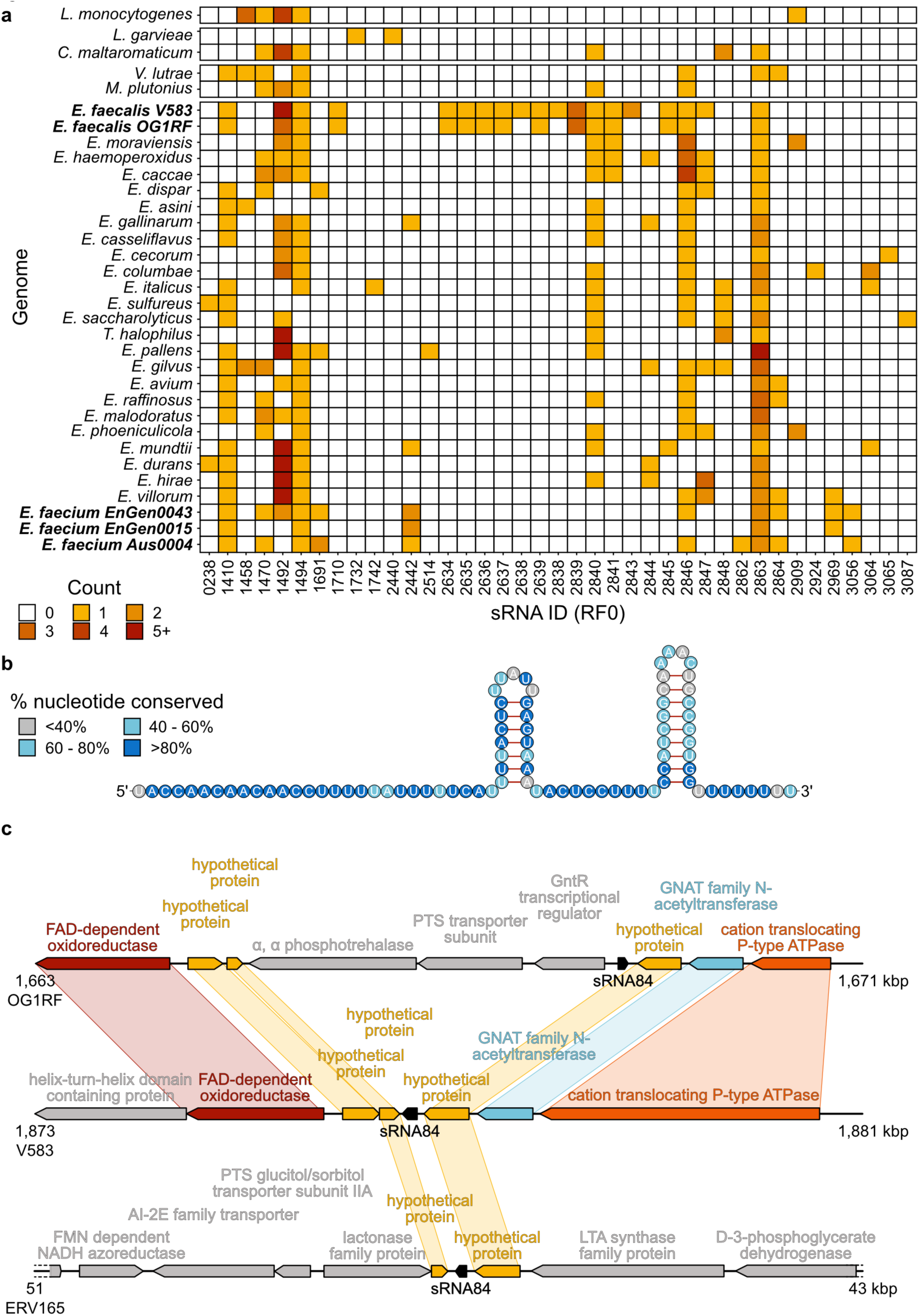
Enterococcus sRNA 84 is a *trans-*encoded sRNA found across the Enterococcaceae. **a)** Heatmap showing presence of sRNAs in genomes across the genus *Enterococcus,* family Enterococcaceae, and other outgroup genomes. Phylogenetic tree adapted from Lebreton et al. 2017. Only RNAs found in at least one *Enterococcus* genome are shown. **b)** Predicted 2-dimensional structure of sRNA 84 with a representative sequence from *E. faecalis* V583 and circles shaded by the degree of conservation across the Enterococcaceae. **c)** Synteny plot of the genetic neighborhood of sRNA 84 in 3 genomes: *E. faecalis* OG1RF (top), *E. faecalis* V583 (middle), and *E. faecium* ERV165 (bottom). Colored genes are found in more than one genome, strain specific genes are shown in grey.

In contrast to these wide-spread sRNAs, there are also three RNAs found in the majority of enterococcal genomes, but not in *L. monocytogenes*: RF02840, RF02846, and RF02863. RF02846 is particularly interesting because it is unique to the Enterococcaceae. RF02846 appears in both pathogenic and commensal strains of *E. faecalis* and *E. faecium*, but not in any genomes outside the family. Given its uniqueness and broad conservation, we wondered if it might regulate genes and functions that are fundamental to enterococci.

RF02846, referred to here as Enterococcus sRNA 84, was first identified in *E. faecalis* by Innocenti et al. in 2015 (Ref84 in strain V583), and in *E. faecium* by Michaux et al. in 2020 (sRNA_101 in AUS0004). While its expression was validated by northern blot (Michaux et al. 2020), its target(s) and function remain unknown. In the *E. faecalis* strain OG1RF, sRNA 84 is predicted to be 86 base pairs long. Secondary structure predictions based on sequence alignment of all members of the family in the Rfam database suggest that the 5’ end of the RNA is single-stranded, followed by two stem-loop structures and a poly-U tail at the 3’ end (Figure 1B). This single-stranded region is highly conserved (Figure S1). Therefore, it may be important for binding to mRNA target sequences. We next examined the genetic neighborhood surrounding sRNA 84 across genomes in different species and strains (Figure 1C). We found that sRNA 84 is intergenic, in many instances between and downstream of two conserved hypothetical genes. The lack of an overlapping mRNA suggests that sRNA 84 is *trans-*encoded, as opposed to an antisense RNA. The locus is highly conserved between *E. faecalis* OG1RF and V583, with the exception of a three gene insertion, which is unique to OG1RF (Bourgogne et al. 2008). By contrast, only the two hypothetical genes (OG1RF_11596; OG1RF_11600) and the sRNA are conserved in *E. faecium* ERV165. Taken together, we find that sRNA 84 is conserved across, as well as specific to, the family Enterococcaceae. With its intergenic location and conserved single stranded binding region, we hypothesize that sRNA 84 may regulate many conserved genes important for enterococci through RNA-RNA interactions.

### sRNA 84 regulates cell wall genes found across the genome

To define sRNA 84’s regulon, we generated a deletion mutant of sRNA 84 in *E. faecalis* OG1RF. We then conducted RNA sequencing of wildtype and ΔsRNA 84 in BHI grown to mid-log phase. In the knockout compared to wildtype, there were 25 differentially expressed genes (Table 1), with 18 of these genes having lower levels in the ΔsRNA 84 strain (Figure 2A), suggesting that sRNA 84 plays a role in activation of these genes in wildtype *E. faecalis*. To understand any possible polar effects due to knocking out sRNA 84, we also sequenced RNA from a knockout strain complemented with sRNA 84 at another location on the chromosome. sRNA 84 expression was similar to that of WT in the complemented strain, with 14 genes differentially expressed (10 upregulated, 4 downregulated) (Figure S2A). Of these, only OG1RF_11599, a GntR transcriptional regulator, was found to be upregulated in both strains (Figure 2C, Figure S2B). This transcription factor is predicted to regulate the downstream PTS system (Bourgogne et al. 2008), similar to other known GntR family members (Tyne et al. 2019). As sRNA 84 is upstream of 11599, editing the sRNA may have impacted 11599 expression. To rule out any downstream impacts of 11599 polar overexpression, we compared RNAseq of a Δ11599 strain and wild type and found no other differentially expressed genes (Figure S2C). Another gene, OG1RF_10328, (LysM domain-containing protein) was significantly downregulated in the knockout, and also significantly upregulated in the complemented strain, possibly due to slight differences in sRNA 84 expression between wildtype and the complement.

**Figure 2:**
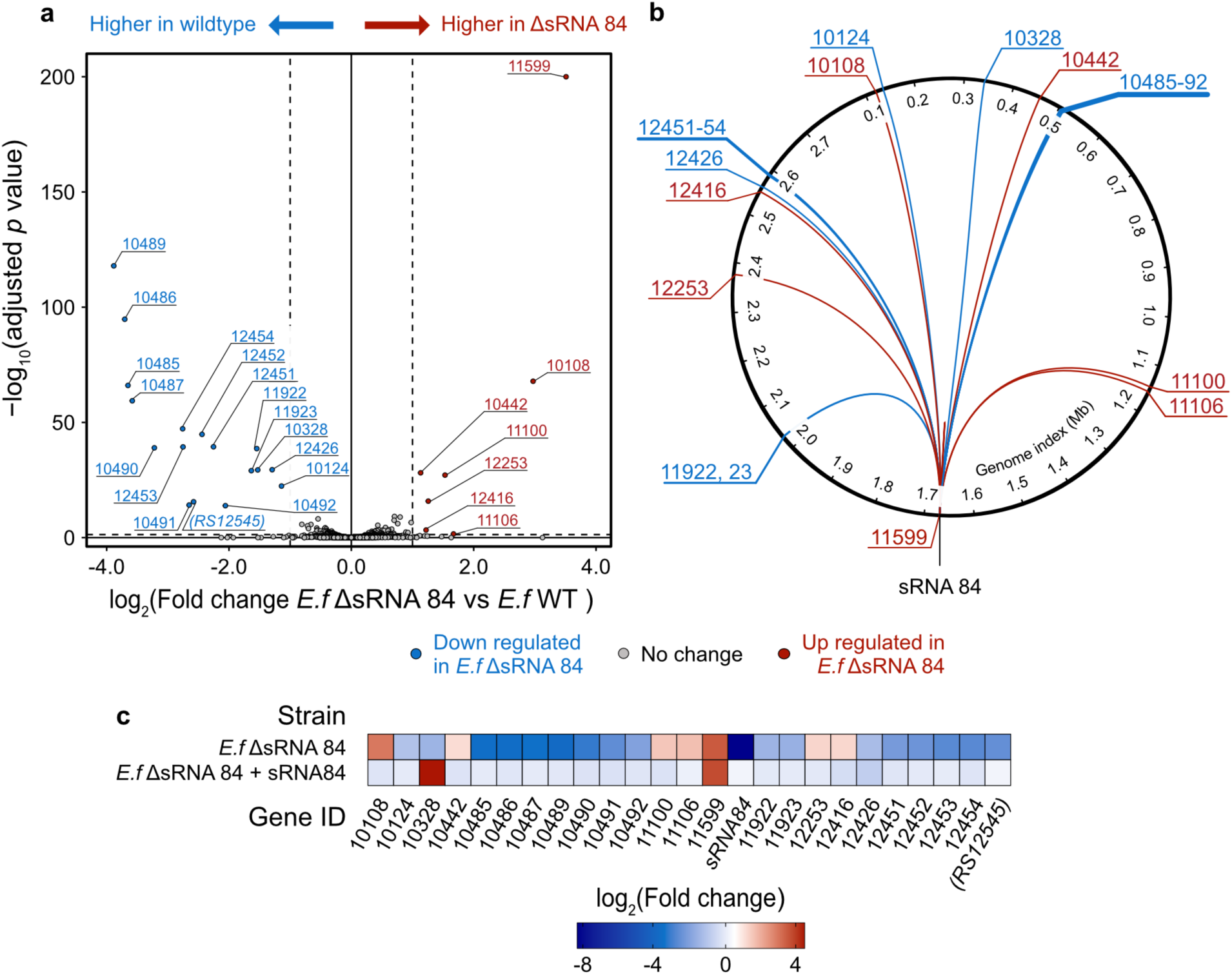
Knockout of sRNA 84 affects expression of cell wall genes across the genome. Genes downregulated in the knockout relative to wild type are in blue. Genes upregulated in the knockout relative to wild type are in red. **a)** Volcano plot of differentially expressed genes (>= 2-fold change, FDR < 0.05, shown as dotted lines) in the mutant vs. wild type. Labels are gene IDs. **b)** Circos plot showing the locations of differentially expressed genes in the genome. Black labels show genetic location in Mb. **c)** Heatmap comparing fold change of differentially expressed genes in the knockout strain vs. wild type to the fold change in the complemented strain vs. wild type. Only genes that are differentially abundant in ΔsRNA 84 are shown.

**Table 1:**
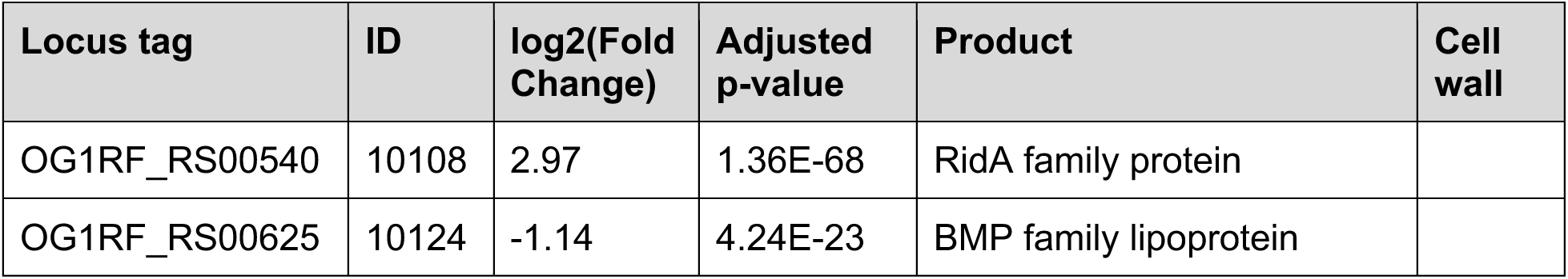
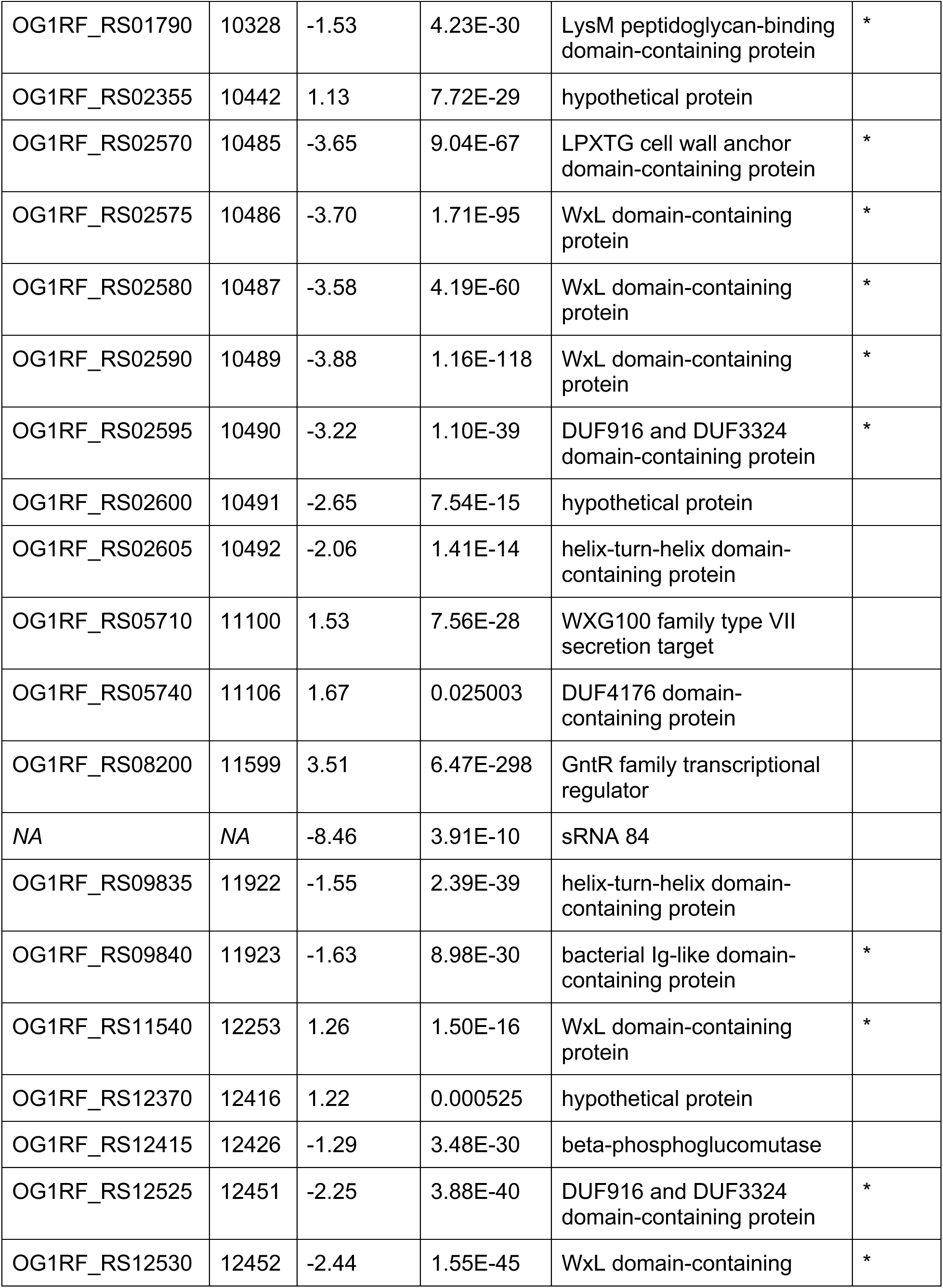
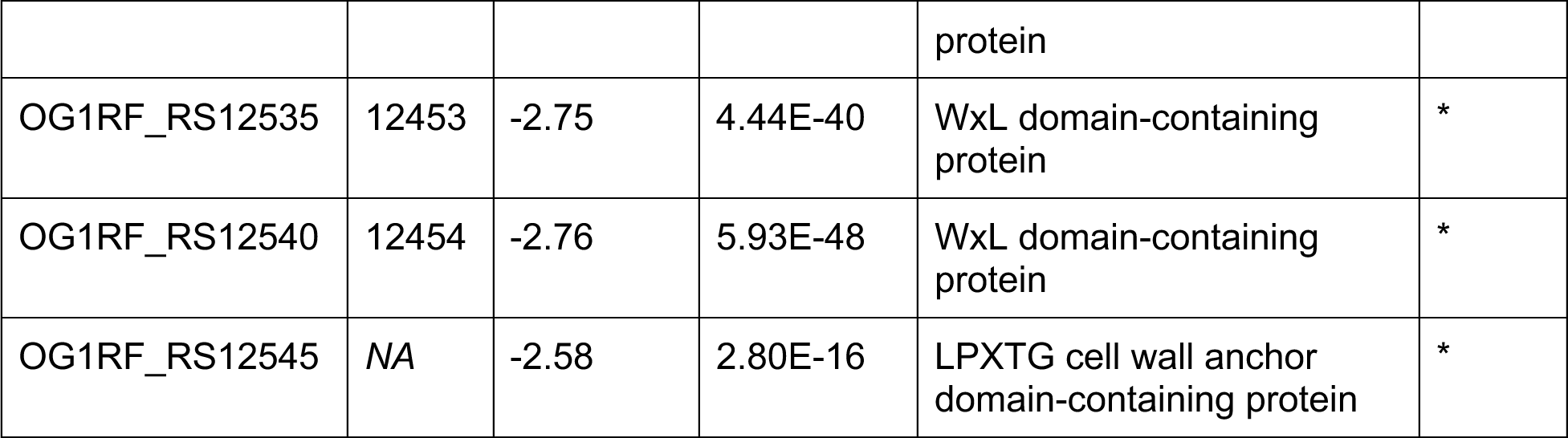
Hits from RNAseq analysis. Sorted by locus ID. An asterisk indicates that this protein may be localized to the cell wall based on domain annotations.

Plotted by their location in the genome, we can see that target genes are distributed across the chromosome—though notably, there are clusters of proximal genes with similar expression patterns (Figure 2B). Upon further inspection, the structure of these regions is consistent with these genes being an operon, implying that they are transcribed as a polycistronic RNA and subject to the same regulation. When examining the annotations of the differentially expressed genes, a pattern emerges: many of the genes that are repressed in the knockout have WxL, LPXTG, or Ig-like domains. These domains are found in proteins localized to the cell wall, many of which are known to play a role in host adhesion and biofilm formation (Galloway-Peña et al 2015; Henrichx et al. 2009). Examining two clusters of differentially expressed cell wall genes (10482 - 10492 & 12448 - 12458), we found arrays of genes with WxL, LPXTG, MucBP (mucin-binding domain), and DUF916/DUF3324 domains. In this experiment, we saw that these genes are all more highly expressed in the wild type vs. the ΔsRNA 84 strain.

Small RNAs can up- or downregulate translation of target genes by either controlling transcript stability or through other mechanisms. To discover if there were additional gene products regulated by sRNA 84 that are not detectable at the RNA level, we analyzed the proteomes of wildtype and ΔsRNA 84 in M9YEG media grown to mid-log phase. Over 13,000 peptides were identified in each sample, with a total of 1,824 proteins detected in at least one sample. Of these, we found 18 proteins were differentially abundant in the knockout compared to wild type (Figure 3A and 3B). Concordant with the RNAseq data, we observed that the WxL-domain containing proteins 10487, 10489, and 12452 were significantly less abundant at the protein level in ΔsRNA 84 than in wild type. Four additional gene products were differentially abundant in the two datasets (Figure 3C and Figure 3D). These were genes 10124 (BMP family lipoprotein), 11599 (GntR transcription factor), 10442 (hypothetical protein), 10108 (RidA family protein). Additionally, some gene products were detected as differentially abundant in the proteomics data but their transcripts were not differentially abundant in the RNAseq data; these include 12396 (ASCH domain-containing protein), 11216 (hypothetical protein), 11384 and 11385 (ABC transporter genes), and 11938 (fumarate reductase). These proteins may be regulated through a mechanism by which the sRNA binding promotes or inhibits translation but does not particularly impact transcript stability.

**Figure 3:**
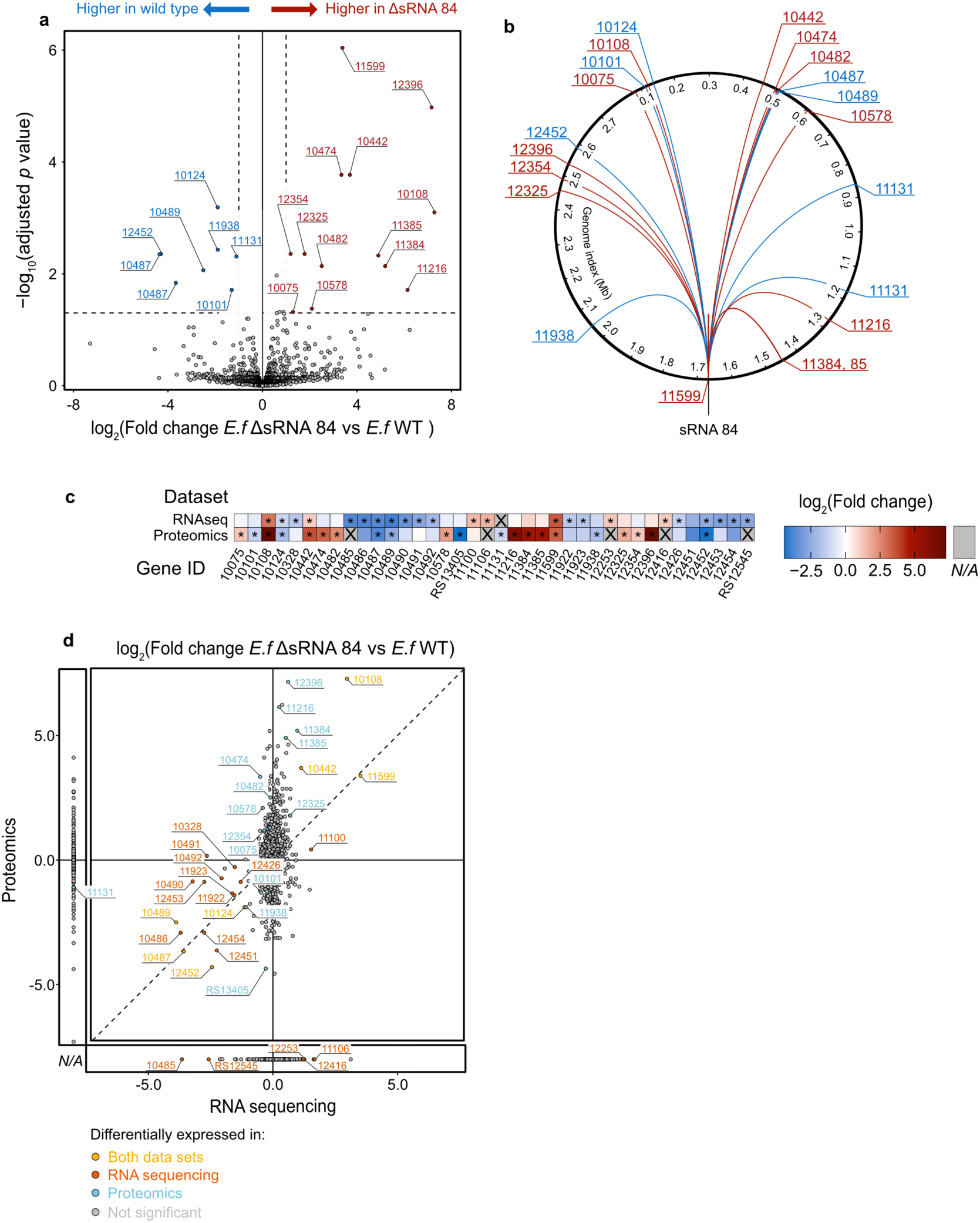
Knockout of sRNA 84 affects expression of genes at the protein level. Genes downregulated in the knockout relative to wild type are in blue. Genes upregulated in the knockout relative to wild type are in red. **a)** Volcano plot of differentially abundant proteins (>= 2-fold change, FDR < 0.05, shown as dotted lines) in the mutant vs. wild type. Labels are gene IDs. **b)** Circos plot showing the locations of differentially abundant proteins in the genome. Black labels show genetic location in Mb. **c)** Heatmap comparing the fold change (knockout relative to wildtype) in the RNAseq vs. proteomics data for any gene found to be differentially expressed in at least one of the datasets. Significant hits (FDR < 0.05) are denoted with an asterisk. Gene products in grey with ‘X’ labels were undetected. **d)** Comparison of fold change (knockout relative to wildtype) in RNA sequencing on the x-axis, and proteomics on the y-axis, colored by significance at alpha = 0.05 (grey - not significant, yellow - significant in both datasets, orange - only in RNAseq, blue - only in proteomics). The dotted line represents where fold changes are equal across the two datasets. Panels on the side of the main plot show genes not detected in a dataset.

sRNA 84 may directly bind to target mRNAs, which may impact their stability and translation. Alternatively, differentially regulated genes may be the result of indirect effects. To assess whether sRNA 84 may base pair with target mRNAs, we used IntaRNA (Mann et al. 2017), to search all mRNAs in the OG1RF genome for predicted interactions with sRNA 84. Indeed, sRNA 84 has sequence complementary to both regions (genes 10485 and 12454, respectively) (Figure 4). This supports the model that sRNA 84 may act as a *trans-*encoded sRNA, base pairing with imperfect complementarity to dozens of targets, and activating expression of many cell wall genes, which may play a role in adhesion to the host.

**Figure 4:**
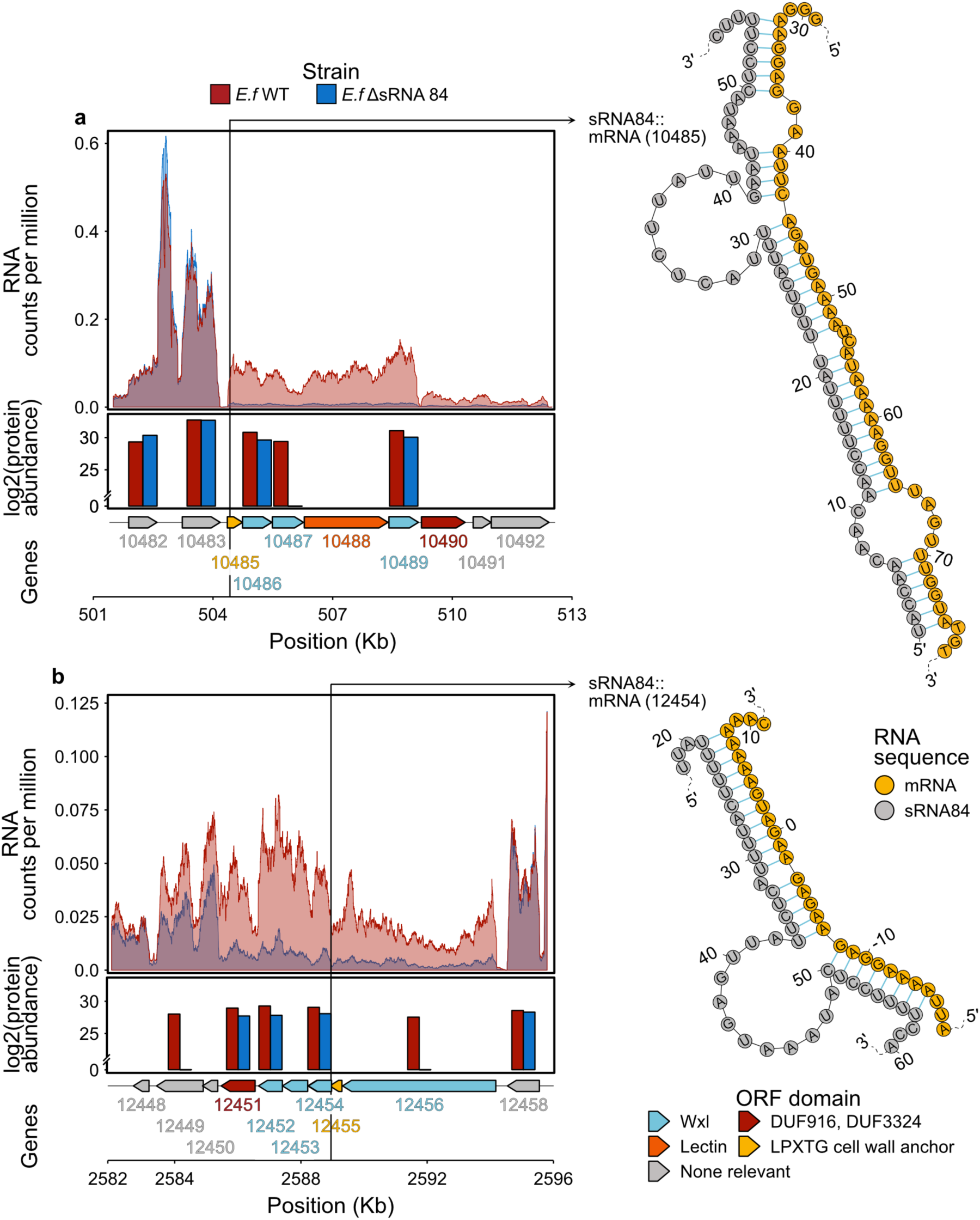
sRNA 84 is predicted to bind to polycistronic mRNAs encoding cell wall genes and may stabilize the transcript. Plot of RNAseq data (mean of 3 replicates, showing counts per million reads at each nucleotide) and proteomics data (mean normalized log_2_(protein abundance)) on the left, with the corresponding operon: genes from 10482 to 10492 **a)**, genes from 12448 to 12458 **b)**. Genes are colored by the presence of relevant cell wall associated domains. The vertical line represents the location of the predicted RNA-RNA interaction with sRNA 84. The IntaRNA predicted interaction is shown on the right; position is relative to the start site for the mRNA.

### sRNA 84 deletion mutant has reduced ability to bind mucin *in vitro*

Given the differential expression of many cell wall proteins, including a putative mucin-binding protein, in ΔsRNA 84, we sought to determine whether sRNA 84 was required for adhesion to mucin *in vitro.* To assess adhesion to mucin, the wildtype strain and the knockout were incubated on mucin-coated plates. Optical density at 600 nm (OD_600_) was read for each culture to ensure no confounding effects due to different cell densities (Figure S3). Cells were then washed off and stained with crystal violet to quantify the remaining bacteria adhered to the plate. Crystal violet stain was normalized to the OD_600_ of each culture. We observed a marked decrease in bacteria bound to mucin in the knockout when compared to wildtype (estimated marginal means 0.378 and 0.493, respectively, p = 4.45e-06) (Figure 5). With reduced expression of many cell wall genes, ΔsRNA 84 is less able to adhere to mucin, suggesting that sRNA 84 may impact the ability of *E. faecalis* to colonize the gastrointestinal tract.

**Figure 5:**
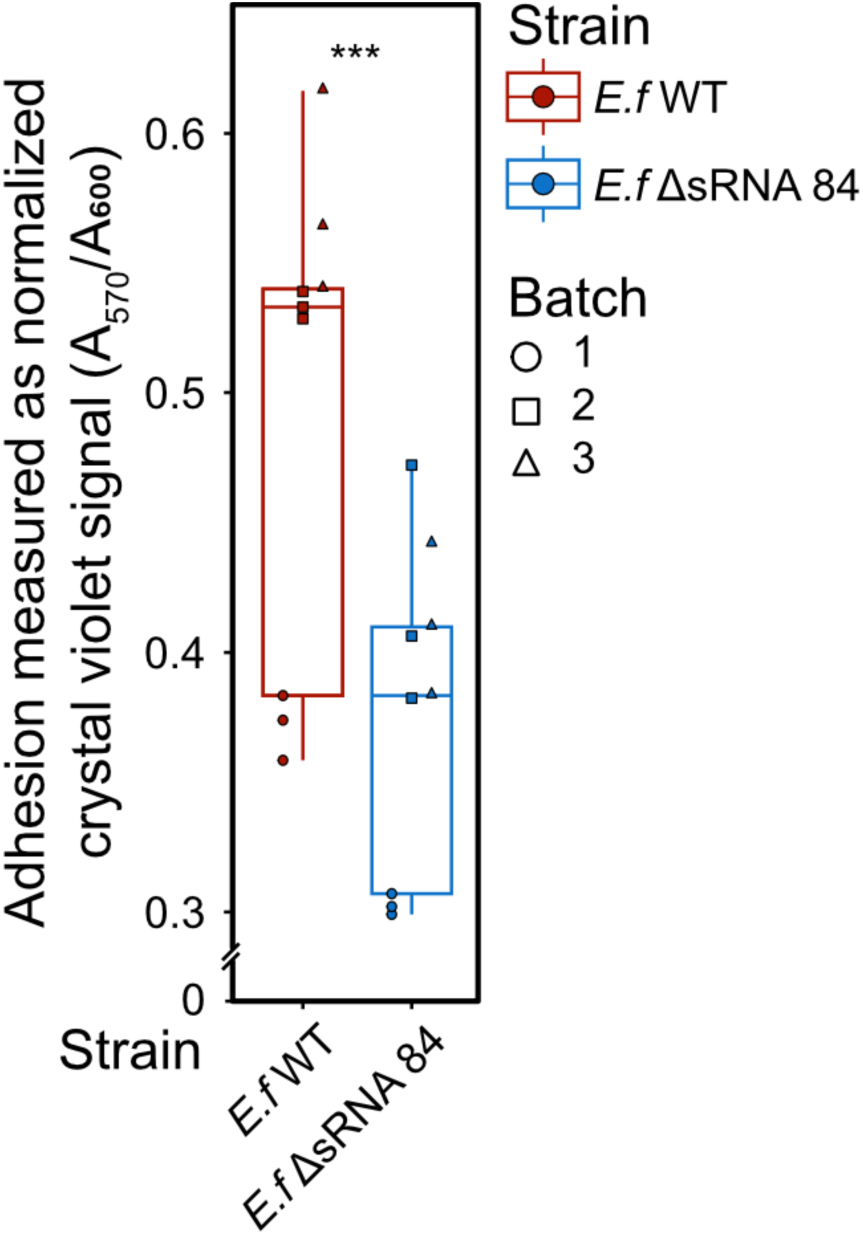
sRNA 84 mutant is deficient in binding mucin *in vitro*. Crystal violet (A_570_) signal normalized to bacterial density (A_600_) of wildtype (red) and the knockout (blue) in the mucin adhesion assay across 3 experiments (shapes) with 3 biological replicates per strain. 2 technical replicates of each biological replicate were averaged. The p-value was calculated using a linear mixed effects model to correct across experimental batches. P-values are indicated as follows: *** p < 0.001.

### sRNA 84 is important for colonization of the gastrointestinal tract

Given the finding that sRNA 84 impacts the ability of *E. faecalis* to bind to mucin, we sought to determine if it also affects colonization of the gastrointestinal tract *in vivo.* To best mimic colonization in a non-dysbiotic setting, we colonized specific-pathogen-free mice whose native microbiomes are unperturbed. This experimental model results in stable long-term colonization of *E. faecalis* in the gastrointestinal tract (Kommineni et al. 2015). During the initial colonization phase, mice were treated with *E. faecalis* in their drinking water for two weeks, receiving either wildtype or the knockout strain. During this time, there were high levels of both wild type and the knockout (Figure 6A). After 14 days, *E. faecalis* treatment ceased and mice were given plain drinking water. In three independent experiments, fecal pellets were collected, diluted, and plated to enumerate colony forming units (CFUs) of *E. faecalis*. One experiment, which did not achieve stable colonization of wildtype, was excluded from analysis (data in Figure S4B). In the remaining batches, both wildtype and the knockout stably colonized the microbiome over the next seven weeks. However, ΔsRNA 84 colonized the gut at approximately 1 log fold lower CFU/g feces compared to the wildtype (Figure 6A). This may be less significant at later timepoints due to the emergence of suppressor mutations, as seen in Banla et al. 2017. On day 63, the mice were sacrificed, and intestinal samples were collected. Colony forming units of *E. faecalis* were enumerated at different positions along the length of the intestine. The ΔsRNA 84 strain was found at lower abundance in the distal small intestine, cecum, colon, and in feces (cecum and feces p-value < 0.05) compared to wild type (Figure 6B). Data for each experimental batch and the native enterococci prior to the start of the experiment is shown in Figure S4.

**Figure 6:**
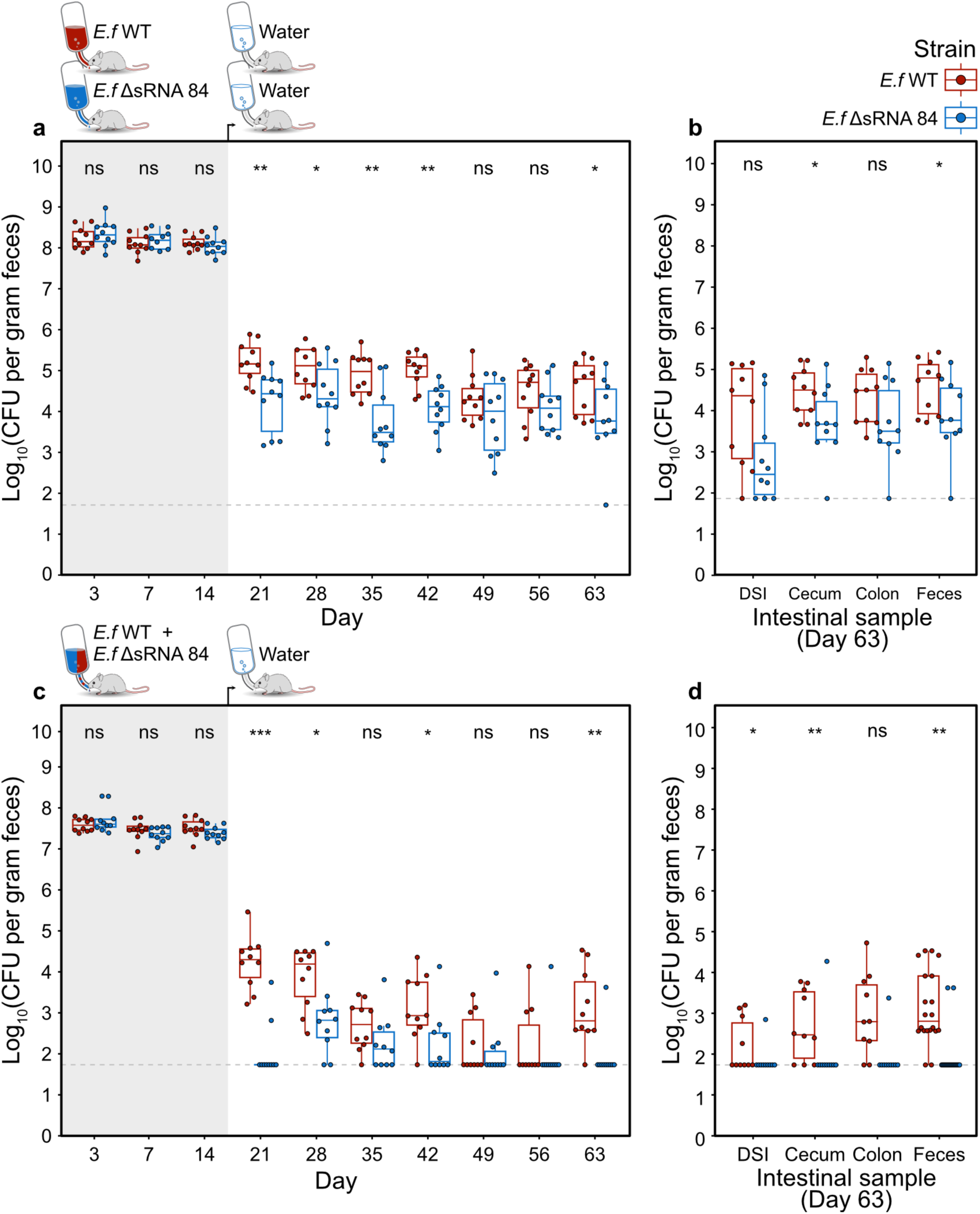
ΔsRNA 84 colonizes the mouse gut at lower abundance and is outcompeted by wildtype. Colonization experiments: mice were given *E. faecalis* in the drinking water starting on day 0 for the 14-day colonization period (grey background), after which they were fed clean drinking water with no bacteria. At each timepoint, feces were collected, diluted, and plated to count CFUs. At the endpoint of the experiment, intestinal samples collected, diluted, and plated to count CFUs. Any plate with less than 10 colonies is below the limit of detection (dashed line representing the lowest CFU/g we detected with at least 10 colonies). To determine statistical significance, a Mann–Whitney U test was conducted at each timepoint across all batches. P-values are indicated as follows: ns p > 0.05, * p < 0.05, ** p < 0.01, *** p < 0.001. The boxplot lines represent the median and upper and lower quartiles, while the whiskers are minima and maxima (not including outliers). **a)** CFU per gram feces of each strain in mice that were separately colonized with either knockout or wildtype *E. faecalis.* N = 5 mice per strain, 2 independent experiments. **b)** CFU per gram of sample (from the distal small intestine (DSI), cecum, colon, and feces) at day 63 from mice which were separately colonized with either knockout or wildtype *E. faecalis.* N = 5 mice per strain, 2 independent experiments. **c)** CFU per gram feces of each antibiotic-labeled strain in mice that were given a 1:1 mixture of wildtype and knockout. N = 10 mice. **d)** CFU per gram of sample (from the distal small intestine (DSI), cecum, colon, and feces) at day 63 from mice which were given a 1:1 mixture of wildtype and knockout. N = 10 mice.

To reduce the impact of mouse-to-mouse variability and assess the impact of sRNA 84 on fitness in a competitive environment, we next conducted a competition assay. Similar to the prior experiment, mice were given *E. faecalis* in their drinking water for two weeks, but this time each mouse received a 1:1 mixture of wildtype and ΔsRNA 84 (labeled with spectinomycin and kanamycin, respectively). During the colonization phase and after removal of *E. faecalis* water, feces were collected, diluted, and plated on BHI + kanamycin or BHI + spectinomycin to count CFUs of both strains. While the *E. faecalis* mixture was being continuously supplied, both strains colonized at equal levels (Figure 6C). However, upon removing a constant source of new bacteria, the wild type outcompeted the knockout (p = 0.00028 at day 21), with very few mice having detectable levels of ΔsRNA 84 at day 21 and at many later timepoints (Figure 6C). Indeed, at the endpoint for this experiment (day 63), ΔsRNA 84 was undetectable in most mice across the different intestinal samples (Figure 6D). To ensure that antibiotic labeling of the strains did not impact their fitness and confound our result, we conducted an additional competition experiment with swapped resistance markers. We again saw significant reduction in CFUs of ΔsRNA 84 recovered from feces at day 21 (Figure S5A) compared to wildtype (Figure S5C), as well as in the distal small intestine, cecum, and colon (S5B), showing that our results are independent of the antibiotic label used for each strain. This suggests that sRNA 84 plays a role in stable colonization of the gastrointestinal tract, especially in a competitive environment— deletion of this sRNA, and therefore reduced expression of cell wall proteins, may make *E. faecalis* less likely to persist in the gut.

## Discussion

Throughout their lifestyles as gut commensals and as infection-causing organisms in wounds, the bloodstream, the urinary tract and beyond, enterococci must adapt their gene expression to survive both in and outside the host. *E. faecalis,* as a noted generalist (Lebreton et al. 2017, Schwartzman et al. 2024), must especially be responsive to changes in the environment. While sRNAs are thought to play key roles in fine tuning and rapidly adapting gene expression in many bacteria, few enterococcal sRNAs have experimentally validated targets or functions. Here, we identified that sRNA 84, which is conserved across and unique to Enterococcaceae, regulates cell wall genes that enable colonization of the gut microbiome through binding to host mucin.

In contrast to many sRNAs which are species- or strain-specific (Livny et al. 2008, Michaux et al. 2020), sRNA 84 is found across the family Enterococcaceae. An intergenic transcript with no coding region and a conserved, single stranded region suggests that sRNA may act as a *trans-* encoded sRNA, regulating many core genes across enterococci. Prior studies defined core genes of special importance to the genus (Gilmore et al. 2020, Lebreton et al. 2017); sRNA 84 joins this list of features conserved across and unique to *Enterococci*. Through RNA sequencing and proteomics analysis of wildtype *E. faecalis* and the ΔsRNA 84 knockout strain, we found that sRNA 84 upregulates dozens of genes, many of them cell wall genes of unknown function. Through computational analysis, we identified predicted interactions between sRNA 84 and these target mRNAs, supporting the model that sRNA 84 functions as a *trans-*encoded sRNA and directly binds to these transcripts and impacts their stability. In addition to hits found via RNAseq, we validated the impact of sRNA 84 on some of these genes at the protein level, and identified additional targets that were only impacted at the translational level. Combined RNA and proteomics data may enable us to learn the rules governing sRNA-mRNA interaction mechanisms in *E. faecalis*.

Given the finding that sRNA 84 upregulated cell wall genes, including genes with structural similarity to known adhesins (Brinster et al. 2006, Hancock et al. 2014, Hendrickx et al. 2009), we sought to assess if sRNA 84 played a role in binding to mucin and colonization of the host. Indeed, we found that ΔsRNA 84 had reduced binding to plates coated in mucin. In a specific-pathogen-free mouse model, ΔsRNA 84 colonized the gut at around a log lower CFU/g feces than wildtype. When put head-to-head in a competition assay, ΔsRNA 84 was outcompeted, demonstrating that sRNA 84 is key in stable colonization of the gut microbiome in a competitive setting. Building on the work of Banla et al. 2019 which identified LPxTG proteins that promote mucin adhesion and colonization, the target genes of sRNA 84, especially WxL genes, may play previously unappreciated roles in the colonization process. To date, no other sRNAs in *E. faecalis* have been found to impact gut microbiome colonization. This expands the known functions of sRNAs in *E. faecalis* beyond virulence and stress response (Michaux et al. 2014).

While we have discovered the targets and function of sRNA 84 in *E. faecalis,* our study has several limitations. First, we predict that sRNA 84 directly binds target mRNAs; however, this was not experimentally confirmed. sRNA-mRNA interaction may also be mediated by unknown proteins that act as chaperones—indeed, the poly-U tail at the 3’ end resembles known Hfq binding sites (Otaka et al. 2011). Additionally, the exact role of the target genes in colonizing the host remain unclear. We don’t know which cell wall proteins bind to mucin, and whether these proteins also bind to other host macromolecules. Lastly, while sRNA 84 is expressed under many laboratory conditions (Michaux et al. 2020), the regulatory cues that influence its expression, especially *in vivo,* remain unknown.

Taken together, this study provides insight into how *E. faecalis* can regulate expression of adhesion proteins to help it persist in the gut microbiome. For a generalist like *E. faecalis*, having a switch to turn on (and off) host-associated genes may help it better adapt to its environment. Given the unique conservation of sRNA 84, understanding the regulation of core genes can provide insight into the mechanisms by which Enterococcaceae interact with hosts and the environment. Whether sRNA 84 plays a similar role in other enterococci, such as *E. faecium,* remains to be seen. In conclusion, sRNA 84, through regulation of adhesins, may contribute to the ubiquity of *E. faecalis* as a commensal organism in the human gut microbiome, a key factor that may drive its prevalence in hospital settings and therefore lead to the high incidence of nosocomial infections. sRNA 84 may also facilitate *E. faecalis* colonization of indwelling catheters, implanted devices, and other host tissues, suggesting it may serve as an exciting target for translational interventions.

## Materials and methods

### Annotation of RNAs in *Enterococci*

24 representative *Enterococcus* genomes, as well as 5 related species were selected based on the phylogeny of enterococci from Lebreton et al. 2017. A *Listeria monocytogenes* outgroup was also added. All RNAs in the RNA families database (RFam) (Ontiveros-Palacios et al. 2025) were annotated in these genomes using cmscan (Nawrocki & Eddy 2013) with the following options: nohhmonly, rfam, cut_ga.

### Sequence and structure conservation analysis

The Stockholm alignment of the Rfam family of Enterococcus sRNA 84 (RF02846) (Ontiveros-Palacios et al. 2025) was visualized in Jalview (Waterhouse et al. 2009). The predicted structure of sRNA 84 was drawn using R2R (Weinberg and Breaker 2011).

### Bacterial culturing

All bacterial cultures, unless otherwise stated, were grown at 37°C and 180 rpm in brain heart infusion (BHI) broth (Millipore Sigma) under aerobic conditions. Antibiotics were used at the following concentrations for *E. faecalis* as required: chloramphenicol 10 µg/mL; kanamycin 500 µg/mL; spectinomycin 100 µg/mL; rifampicin 100 µg/mL. M9YEG media was made with: M9 salts supplemented with yeast extract (0.25%) and glucose.

### Mutant construction

The knockout of sRNA 84 was constructed using the homologous recombination system described in Kristich et al. 2007, Vesic and Kristich 2014, Kellogg et al. 2017, and Zlitni et al. 2024. PCR fragments of the pJH086 backbone (provided by the Kristich lab) and 1 Kb fragments up- and downstream of the desired deletion were generated using Q5 mastermix (New England Biolabs) and assembled using NEBuilder HiFi Assembly. Assemblies were transformed into NEB Beta-10 cells, then grown, miniprepped (Qiagen), and sequence verified by nanopore sequencing (Plasmidsaurus). Constructs were then transformed into competent *E. faecalis* via electroporation using a BioRad GenePulser on setting EC1. Competent cells were prepared using the glycine method (Cruz-Rodz and Gilmore 1990). After electroporation, 1 mL of BHI was immediately added to the cells, then incubated on ice for 5 minutes, followed by 90 minutes of outgrowth at 30°C. Transformations were plated on selection plates (BHI containing chloramphenicol and 150 µg/mL X-gal) and grown 1-2 days at 30°C. Transformants were re-streaked for single colonies. Then, integrants were selected by culturing at 42°C on selection plates. After re-streaking for single colonies, integrants were streaked on counterselection plates (MM9YEG agar with 10mM p-Cl-Phe and 150 µg/mL X-gal) at 30°C. Colonies were patched on plates with chloramphenicol to confirm the loss of the plasmid, then screened for mutations at the desired site using colony PCR. All mutants were confirmed using whole-genome nanopore sequencing (Plasmidsaurus). Complemented and labeled strains had the gene of interest introduced between OG1RF_11778 and OG1RF_11779, a safe site for knock-ins as described in DebRoy et al. 2012.

### RNA sequencing

RNA extraction and sequencing analysis were conducted as described previously (Zlitni et al. 2024). Briefly, 3 overnight cultures of each strain were set up in BHI, then subcultured 1/100 for 3 hours to OD_600_ ∼0.7. 4 mL of culture was then quenched with 0.5 mL 9:1 ethanol:phenol. Cells were pelleted, resuspended in 250 µL PBS, and then lysed by treating with 10 µL lysozyme (Millipore Sigma L3790) for 30 minutes at 37°C, followed by treatment with 30 µL of 20% SDS for 30 minutes at 37°C. After lysis, 1.5 mL of TRIzol was added. Following a 10 minute incubation on the bench, 0.5 mL of chloroform was added to each sample and mixed via vortexing. After a 3 minute incubation on the bench, samples were centrifuged at 13,000 x g for 10 minutes. The aqueous phase was then removed, and RNA was extracted according to manufacturer’s instructions using the Zymo RNA Clean & Concentrator-5 kit. rRNA depletion (Novogene-tailored Illumina RiboZero Plus), library preparation (paired-end 150 bp stranded), and sequencing (Illumina NovaSeq 6000 platform) were conducted by Novogene.

For RNA sequencing analysis: the reads were trimmed and filtered using trim galore (Krueger 2015), mapped to the OG1RF reference genome using bowtie2 (Langmead et al. 2009), and coverage of each gene was counted using bedtools coverage (Quinlan and Hall 2010). Differential expression analysis was conducted using DeSeq2 (Love et al. 2014).

### Proteomic Sample Preparation, LC–MS/MS Analysis, and Data Processing

Three replicates of each strain were grown overnight in BHI, then each replicate was subcultured 1/100 in M9YEG and grown for 4 hours to an OD ∼0.5. Cultures were pelleted, washed twice with cold 10 mM tris-HCl pH 8, and snap frozen. Samples were then shipped on dry ice and stored at -80°C until used. Sample lysis, protein extraction, and mass spectrometry were conducted as detailed in Davin et al. 2024. Samples were thawed on ice, then bead-beaten using a Cryogrinder at 1750 rpm for 5 minutes to penetrate cell membranes. For lysis, samples were treated with 4% SDS and 10 mM dithiothreitol, and then heat treated at 90 °C for 10 minutes. Proteins from 300 µg of lysed samples were extracted using a protein aggregation capture (PAC) approach (Batth et al. 2019) and digested with trypsin at a protein-to-enzyme ratio of 1:75. The digested peptide mixtures were analyzed on a Vanquish UHPLC system coupled to an Orbitrap Q-Exactive Plus mass spectrometer (Thermo Fisher Scientific) using a one-dimensional liquid chromatography–tandem mass spectrometry (LC–MS/MS) workflow. 3 µg of peptides per sample were loaded onto a 5 µm Kinetex Phenomenex trapping column prior to separation on a 1.7 µm Kinetex Phenomenex analytical column (made in-house). Desalting was performed for 30 minutes using solvent A (0.1% formic acid in 5% acetonitrile). Peptide separation and elution were carried out over 3 hours with a linear gradient increasing from 0 to 30% solvent B (0.1% formic acid in 70% acetonitrile), followed by a column wash. Data-dependent acquisition was used. Full MS scans were collected across an m/z range of 300– 1,500 at 70,000 resolution (full width at half-maximum). The 20 most intense precursor ions were selected for MS/MS fragmentation using a 1.8 m/z isolation window and a dynamic exclusion of 30 seconds. Peptide identification was performed with Proteome Discoverer (v3.2.0, Thermo Fisher Scientific). Spectra were searched against the *E. faecalis* CDS database (from RefSeq, annotated with PGAP) as well as a common contaminants database using a target-decoy strategy. Search parameters included static cysteine carbamidomethylation and dynamic methionine oxidation modifications, fully tryptic peptides, allowance for up to two missed cleavages, and a minimum peptide length of seven amino acids. Match between runs were not enabled, and a 1% false discovery rate threshold at the peptide level was applied. Peptide quantification was based on area under the curve, and protein-level abundances were calculated by summing the intensities of unique peptides assigned to each protein. Protein abundance was normalized by log_2_ transformation, mean centering, and Lowess normalization. Missing values were imputed. Fold changes and p-values were determined using limma (Ritchie et al. 2015) with Benjamini-Hochberg multiple hypothesis correction. Thresholds for significance were |log_2_(fold change)| > 1 and adjusted p-value < 0.05.

### RNA-RNA interaction predictions

IntaRNA (Mann et al. 2017, Busch et al. 2008, Raden et al. 2018, Wright et al. 2014) was used to predict interactions between sRNA 84 (sequence from OG1RF) against all possible target RNA sequences in the OG1RF NCBI reference genome using default parameters. All base pair positions are reported relative to the start site for each target.

### *In vitro* mucin adhesion assay

The mucin adhesion assay was adapted from Banla et al. 2019. 24-well tissue culture plates were incubated with 0.25% mucin (from porcine stomach, Type III, Sigma-Aldrich) overnight at 4°C. 3 biological replicates of each strain were grown as overnight cultures in BHI. The next day, mucin was removed and the plate washed twice with PBS. The strains were subcultured 1/50 for 2 hours to OD ∼0.6. Cultures were then spun, washed with twice PBS, and resuspended to the same OD_600_. Cells were incubated on the plate for 1 hour at 37°C—for each biological replicate, 2 wells were used as technical replicates. Control wells received only PBS. OD_600_ of each culture was measured using the Synergy H1 plate reader. Then cells were removed, wells were washed twice with PBS and then strained with crystal violet. Following incubation in the dark for 30 minutes, the stain was removed, the wells were washed twice with PBS, and then left to dry. The stain was solubilized with 80:20 ethanol:acetone and read on the plate reader in absorbance mode at 570 nm. Crystal violet signal was normalized to OD_600_ of each culture. P-values were calculated using lmerTest (Kuznetsova et al. 2017) to correct for differences between experimental batches and estimated marginal means for each group were generated with emmeans (Lenth & Piaskowski 2025).

### Animal protocols

All protocols have been approved by the committee for animal care and use at the Medical College of Wisconsin. Five-to six-week-old male C57BL/6J mice were obtained from the Jackson Laboratory (room RB08, JAXWEST facility, Sacramento, CA). Mice were acclimatized for 1 week prior to conducting experiments. Mice were fed a standard chow diet (PicoLab laboratory rodent diet) and de-chlorinated reverse osmosis (RO) water ad libitum. All mice were maintained under specific-pathogen-free conditions throughout the course of the experiments and mice that were colonized by experimental *E. faecalis* strains were housed in an ABSL2 facility. All mice were humanely euthanized by CO_2_ asphyxiation followed by cervical dislocation.

### *E. faecalis* colonization and quantification

Cultures of *E. faecalis* were grown to stationary phase overnight in Mueller-Hinton broth (Difco), supplemented with 50 µg/mL rifampicin (Chem-impex), at 37°C and shaking at 200 rpm. Cultures were pelleted by centrifugation at 4,000 rcf for 10 minutes and the pellets were washed with autoclaved Milli-Q water twice to remove excess media and antibiotics. Final pellets were resuspended in autoclaved Milli-Q water, and bacterial cell concentration was determined by measuring the OD_600_ with a spectrophotometer (NanoDrop 2000 spectrophotometer; Thermo Scientific). Colonization was performed as described previously (Kommineni et al. 2015, Banla et al. 2019). At day 0, native enterococcal load was enumerated by plating feces on selective m Enterococcus (mEnt) agar. Then, mice were fed 5 × 10^8^ CFU/mL of bacteria in 300 mL of drinking water which was replaced every 3-4 days. *E. faecalis*-containing water was removed and replaced by sterile drinking water 14 days following the start of feeding. Fresh fecal samples or representative intestinal samples were weighed and serial dilutions were plated on BHI agar (BD Biosciences), supplemented with 200 µg/mL rifampicin, and incubated at 37°C overnight to enumerate *E. faecalis* CFU/g sample.

For competition experiments, cultures of each strain were pelleted, washed, and resuspended at 5 × 10^8^ CFU/mL. Equal parts were combined to generate 200mL of 5 × 10^8^ CFU/mL water. At day 0, native enterococcal load was enumerated by plating feces on mEnt agar. At each subsequent timepoint, CFUs of each strain were determined by diluting feces and plating on BHI agar supplemented with 500 µg/mL kanamycin (VWR) for the WT (strain EF136) or 100 µg/mL spectinomycin (TCI America) for the mutant (EF155).

## Supporting information

Supplemental File 1 - Strains

Supplement File 2 - Primers

Supplemental File 3 - Plasmids

Supplemental File 4 - sRNA counts

Supplemental File 5 - MSA

Supplemental File 6 - RNAseq

Supplemental File 7 - Proteomics

Supplemental File 8 - IntaRNA

Supplemental File 9 - Adhesion

Supplemental File 10 - Colonization

Supplemental File 11 - Competition

## Data availability

RNA sequencing data generated in this study is available on SRA under BioProject ID PRJNA1348591. Proteomics data is available on ProteomeXChange under the MassIVE dataset ID MSV000099573.

## Author contributions

A.S.B, S.B, and B.J.F. conceived of the study. S.B, S.Z, and E.F.B. generated mutant strains and extracted RNA samples. Proteomics experiments were designed by S.B, M.E.D, P.W, R.L.H, and A.S.B, with the experiment and analysis conducted by S.B. and M.E.D. Mouse experiments were designed by S.B, K.C.J, N.H.S, and A.S.B and conducted by K.C.J. and L.A. Computational analysis was performed by S.B. The figures were drafted by S.B. and A.N. The manuscript was drafted by S.B, and finalized by S.B, A.S.B, and A.N. All authors read and approved the final manuscript.

## Acknowledgements

We thank members of the Bhatt lab for their helpful suggestions and feedback. We thank Dr. Chris Kristich, Dr. James Collins, Dr. Michelle Chua, Dr. Howard Hang, and Dr. Victor Chen for sharing reagents and expertise for generating deletion mutants. We are grateful to Dr. Denise Monack and Ramya Narasimhan for their guidance in developing mouse colonization protocols. We thank the Stanford Genome Technology Center, Fernando Campo and Dr. Laurel Crosby for the use of the Omnilog instrument. This work was supported by a National Institutes of Health R01 AI148623 (A.S.B.), National Institutes of Health R01 AI143757 (A.S.B.), Paul Allen Foundation Distinguished Investigator Award (A.S.B.), Stand Up to Cancer Convergence Award (A.S.B.) and a Center for Pediatric IBD and Celiac Disease Seed Grant (309906). This work was also supported by a Fulbright Research Scholarship from the Commission for Educational Exchange between the United States, Belgium and Luxembourg to P.W. Additional funding was provided by the Luxembourg National Research Fund under INTERMOBILITY/23/17856242.

## Supplementary Figures

**Supplemental Figure 1:**
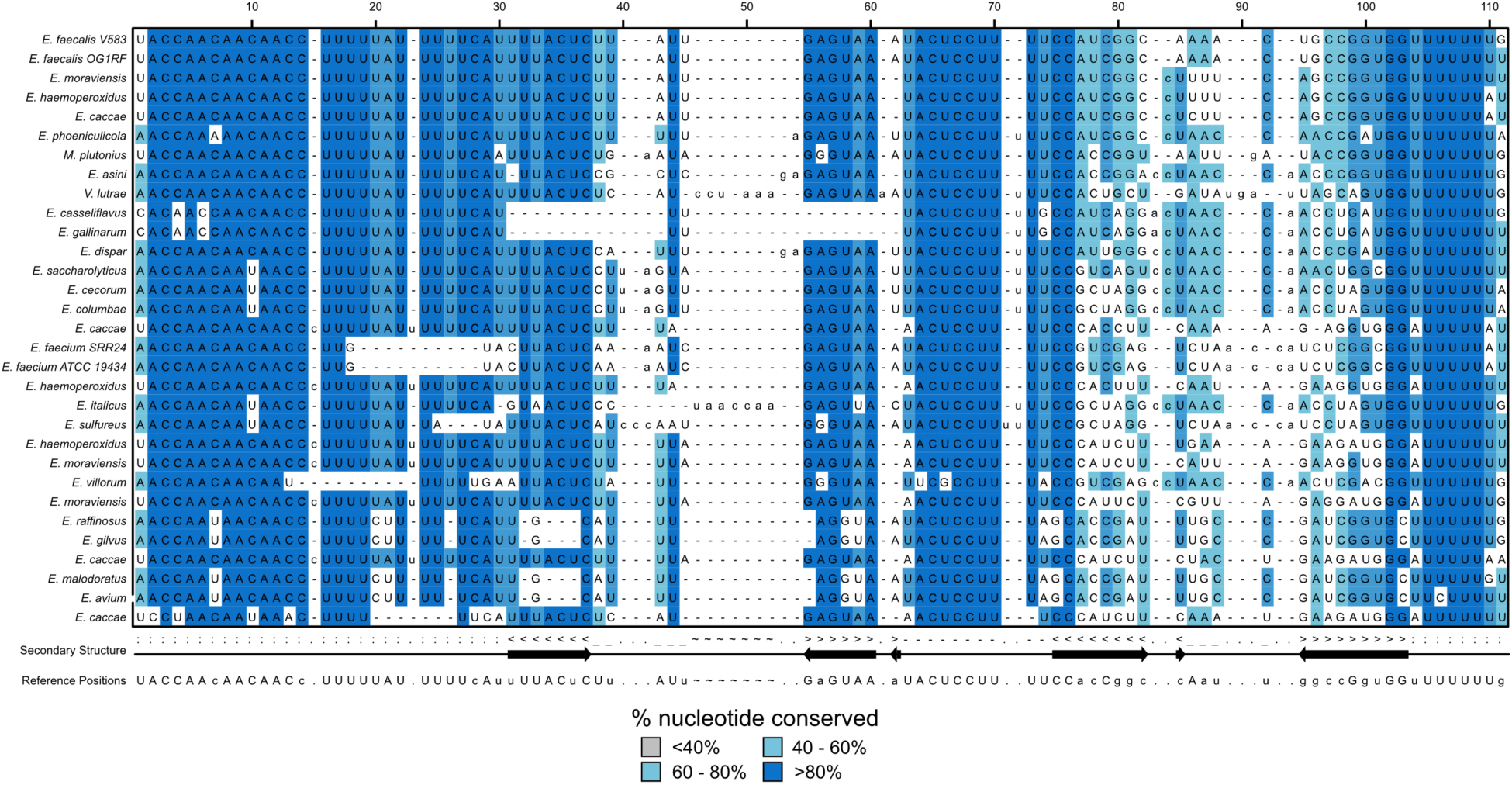
Conservation of sRNA 84 sequence and structure. Multiple sequence alignment of members of the RF02846 family from the RNA families (RFam) database. Blue nucleotides represent the consensus base at that position, with darker color being more highly conserved. Consensus sequence and structure are detailed at the bottom.

**Supplemental Figure 2:**
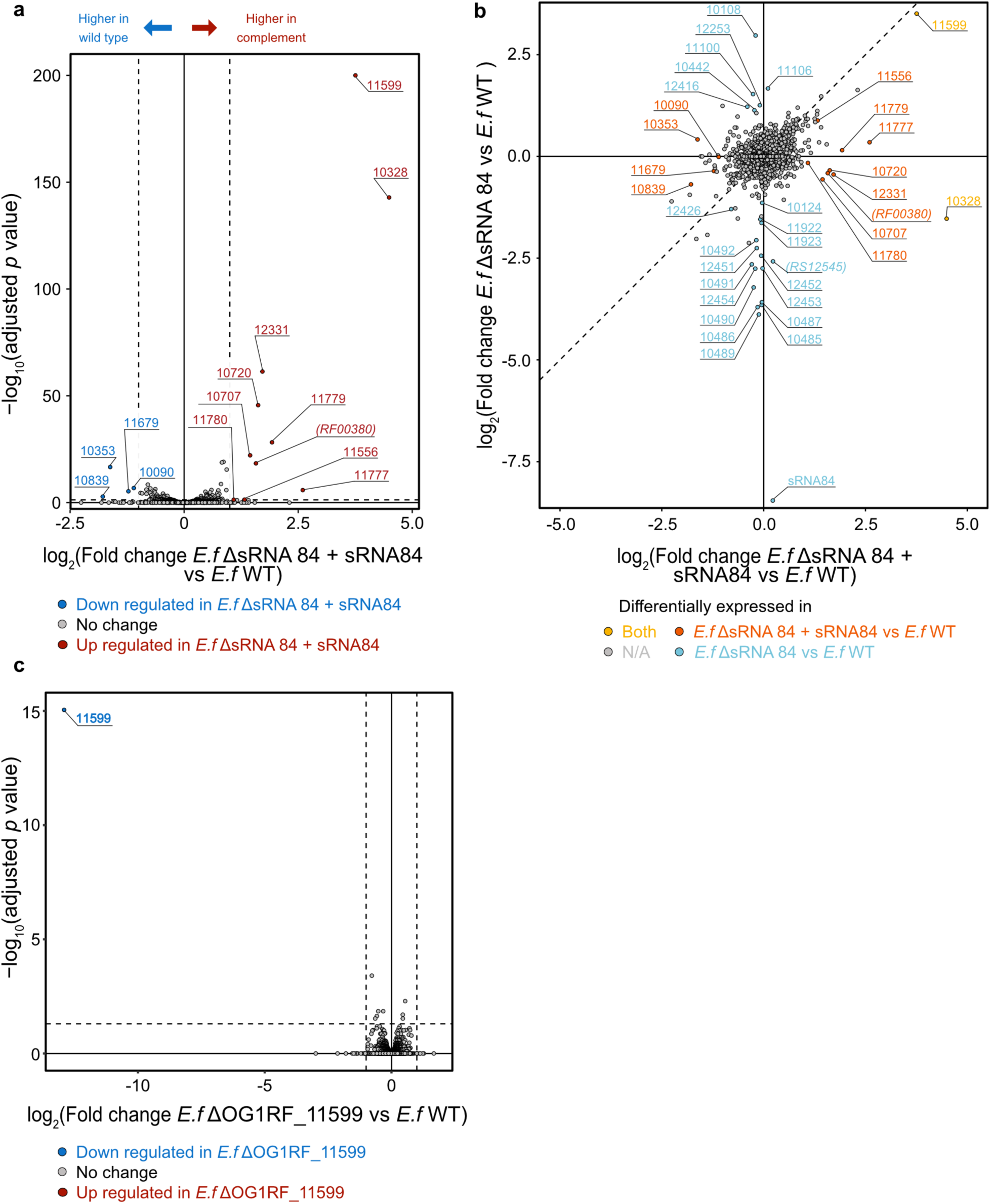
RNAseq of complemented strain. **a)** Volcano plot of complemented strain vs WT. P-values are capped at -log10(200). Dotted lines represent thresholds for significance: p = 0.05, |log2(fold change)| > 1. **b)** Comparison of fold change in knockout vs. WT, and complemented vs. WT, colored by significance at alpha = 0.05 (grey - not significant, yellow - significant in both datasets, orange - only in complement, blue - only in knockout). The dotted line represents where fold change in the knockout is equal to the fold change in the complemented strain. **c)** Volcano plot of GntR transcription factor knockout (ΔOG1RF_11599) vs WT. Dotted lines represent thresholds for significance: p = 0.05, |log2(fold change)| > 1.

**Supplemental Figure 3:**
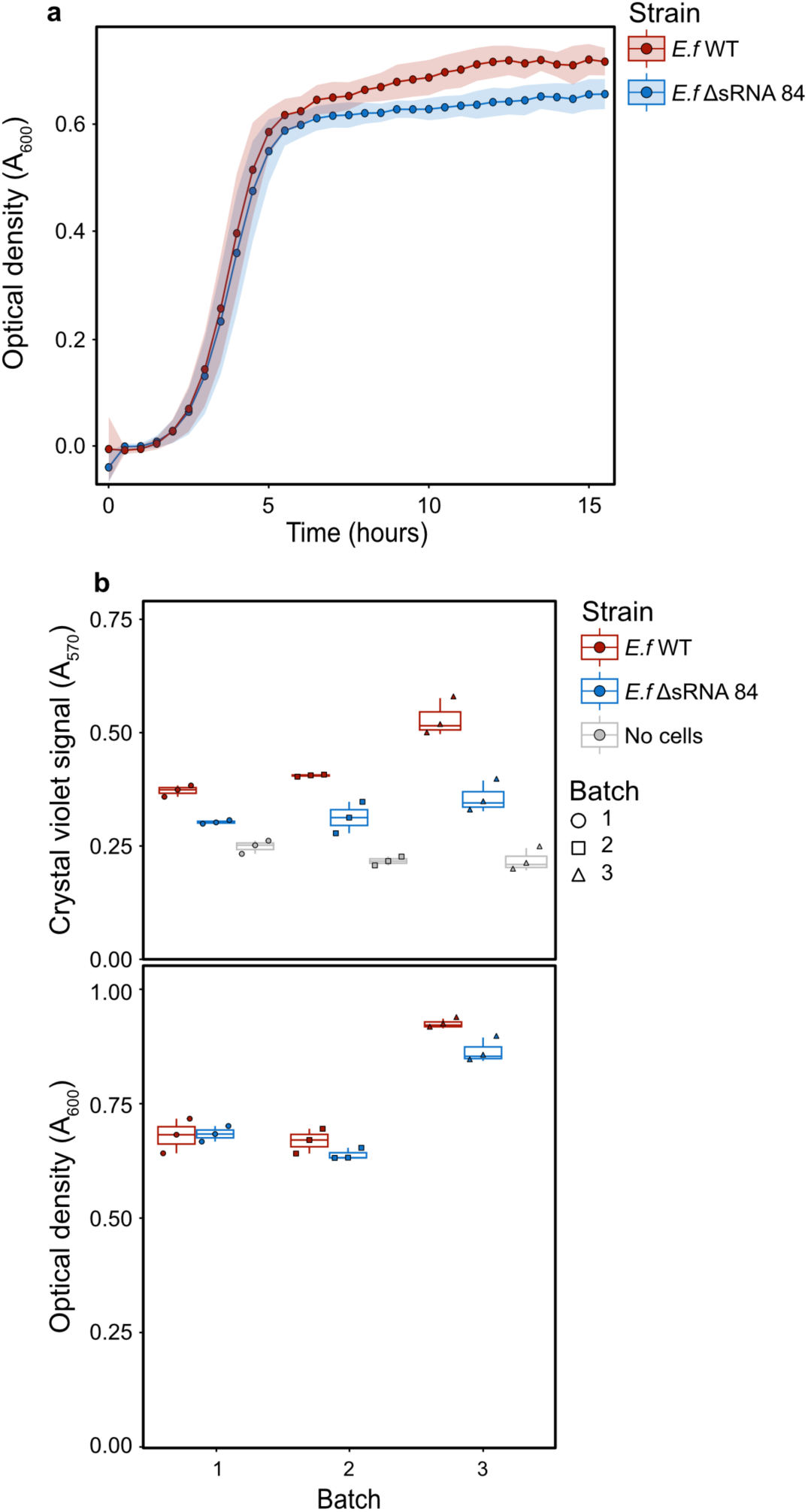
Growth and mucin binding properties of sRNA 84 knockout *E. faecalis* compared to wildtype. **a)** Growth curve data of wildtype *E. faecalis* and the ΔsRNA 84 mutant. **b)** Raw crystal violet (A_570_) and cell density (A_600_) data from the mucin adhesion experiment for each batch of wildtype and ΔsRNA 84, including no cell controls.

**Supplemental Figure 4:**
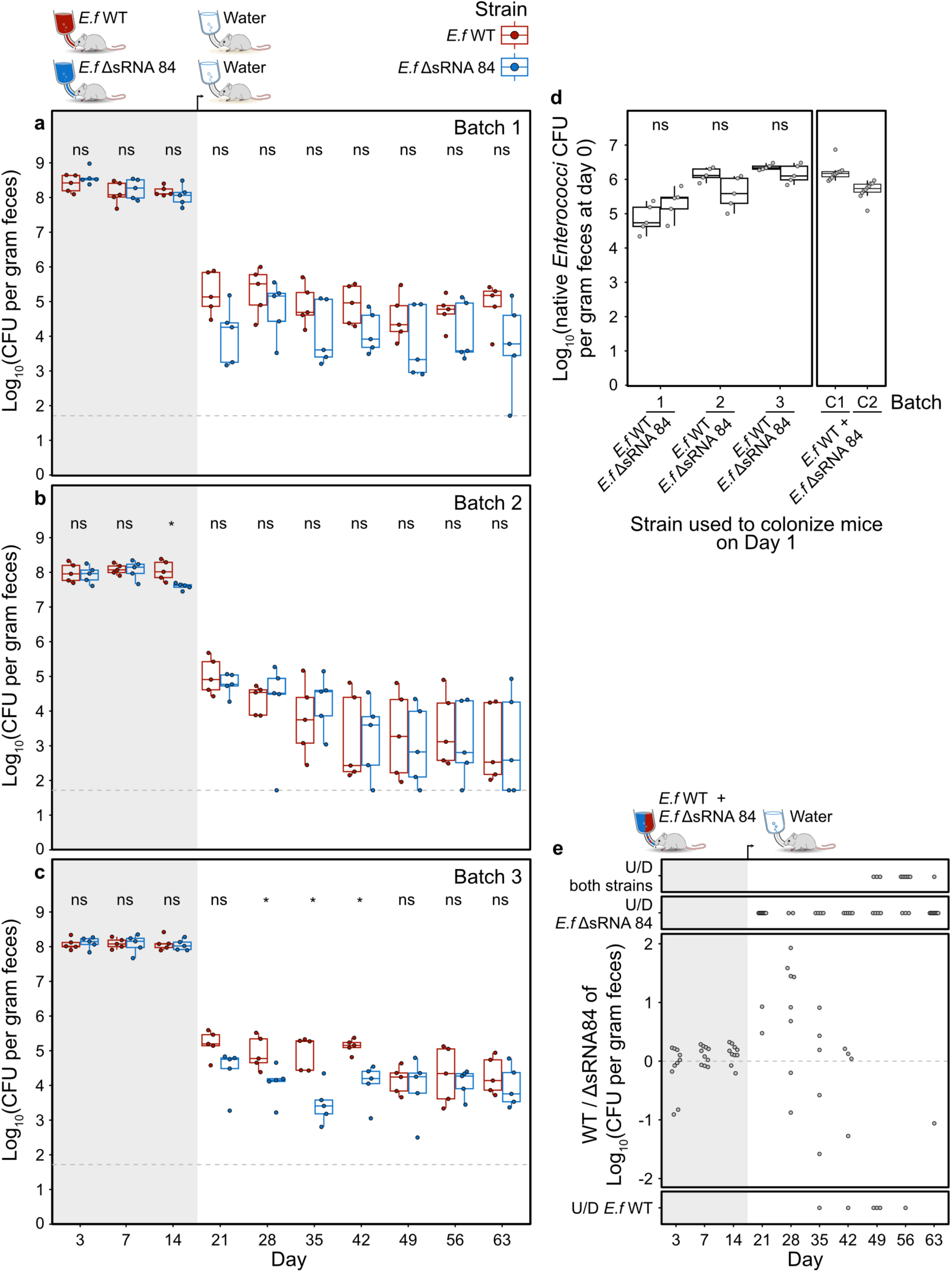
Colonization experiment data for each separate experimental batch. Mice were given either knockout or the wild type strain in the drinking water starting on day 0 for the 14-day colonization period (grey background), after which they were fed clean drinking water with no bacteria. At each timepoint, feces were collected, diluted, and plated to count CFUs. Any plate with less than 10 colonies is below the limit of detection (dashed line representing the lowest CFU/g we detected with at least 10 colonies). To determine statistical significance, a Mann–Whitney U test was conducted at each timepoint across all batches. P-values are indicated as follows: ns p > 0.05, * p < 0.05. The boxplot lines represent the median and upper and lower quartiles, while the whiskers are minima and maxima (not including outliers). The experiment was conducted as 3 experiments with 5 mice each. The data from each replicate is plotted independently: **a)** Batch 1, **b)** Batch 2, **c)** Batch 3. **d)** Native enterococci in each experimental group before the start of the experiment for the colonization experiments (1,2,3) and competition experiments (C1,C2). **e)** Competitive index for competition experiment. The mice where one or both strains were below the limit of detection are marked as undefined above and below the main plot.

**Supplemental Figure 5:**
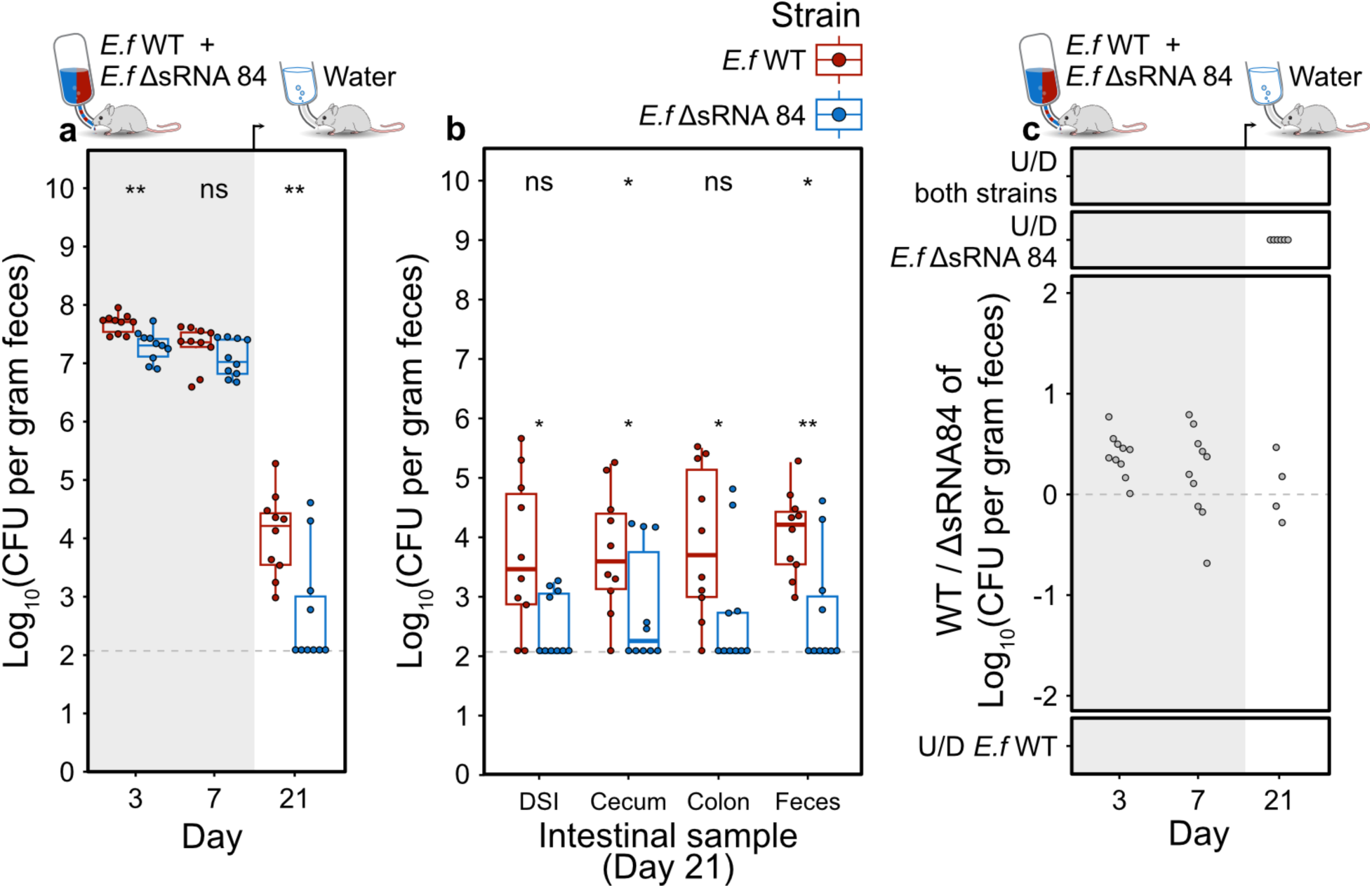
Competition experiment with swapped resistance markers. Mice were given a 1:1 mixture of knockout and wild type strains in the drinking water starting on day 0 for the 14-day colonization period (grey background), after which they were fed clean drinking water with no bacteria. At each timepoint, feces were collected, diluted, and plated to count CFUs. Any plate with less than 10 colonies is below the limit of detection (dashed line representing the lowest CFU/g we detected with at least 10 colonies). To determine statistical significance, a Mann–Whitney U test was conducted at each timepoint across all batches. P-values are indicated as follows: ns p > 0.05, * p < 0.05. The boxplot lines represent the median and upper and lower quartiles, while the whiskers are minima and maxima (not including outliers). **a)** CFU per gram feces of WT and knockout at days 3, 7, and 21. **b)** CFU per gram of each intestinal sample at the endpoint of the experiment, day 21. DSI = distal small intestine. **c)** Competitive index of WT / knockout. The mice where one or both strains were below the limit of detection are marked as undefined above and below the main plot.

